# Neural correlates of newsvendor-based decision making in the human brain: *An exploratory study to link neuroeconomics with neuroimaging using fNIRS*

**DOI:** 10.1101/2020.02.08.940197

**Authors:** Hashini Wanniarachchi, Yan Lang, Xinlong Wang, Sridhar Nerur, Kay-Yut Chen, Hanli Liu

**Affiliations:** Department of Bioengineering, University of Texas at Arlington, Arlington, TX 76019; Department of Information Systems and Operations Management, University of Texas at Arlington, Arlington, TX 76019

## Abstract

Neuroeconomics with neuroimaging is a novel approach involving economics and neuroscience. The newsvendor problem (NP) is a prevalent economics concept that may be used to map brain activations during NP-evoked risky decision making. In this study, we hypothesized that key brain regions responsible for NP are dorsolateral prefrontal cortex (DLPFC) and orbitofrontal cortex (OFC). Twenty-seven human subjects participated in the study using 40 NP trials; the participants were randomly assigned to a group with a low-profit margin (LM) or high-profit margin (HM) treatment. Cerebral hemodynamic responses were recorded simultaneously during the NP experiments from all participants with a 77-channel functional Near-infrared Spectroscopy (fNIRS) system. After data preprocessing, general linear model was applied to generate brain activation maps, followed by statistical t-tests. The results showed that: (a) DLPFC and OFC were significantly evoked by NP versus baseline regardless of treatment types; (b) DLPFC and OFC were activated by HM versus baseline; and (c) DLPFC was activated during LM versus baseline. Furthermore, significant deactivation in right DLPFC was shown due to LM with respect to HM. This study affirms that DLPFC and OFC are two key cortical regions when solving NP. In particular, right DLPFC was found to be more deactivated under challenging risk decision making.

## Introduction

Neuroeconomics is an emerging field integrating economic theories with neuroscience to enhance the understanding of how humans make decisions under different cost-effective conditions^1^. The “behavioral perspective in decision making” is common in business research. For example, studies in management science have found that psychological factors, particularly attitudes towards risks and rewards, are main drivers of business decisions. In general, business research usually assesses cognitive and psychological processes indirectly via survey and/or observations of behavior. The current, largely behavioral-focused, literature on inventory decision making relies on “guessing” the underlying cognitive or psychological mechanism from observations, without any direct physical and/or neurological evidence. On the other hand, functional neuroimaging is a unique tool that allows researchers to “peek into the black box” and gather data directly from pertinent regions of the brain while subjects are engaged in making decision. Advanced neuroimaging technologies, such as multi-channel electroencephalography (EEG) and functional magnetic resonance imaging (fMRI), are potentially able to provide clues to the underlying cognitive processes that hitherto were assumed to be the determinants of human decision-making.

However, neuroeconomics with neuroimaging is still in its infancy because of many challenges ^2,3^. For example, the fMRI measurement settings are very different from naturalistic environments, limiting accurate explanatory power for real-life decision-making processes. Also, it is uncertain “whether neuroimaging can provide theories for economists or whether economic theories can provide frameworks for neuroscience”^2^. Thus, in this study, we wish to explore a non-fMRI neuroimaging approach, namely multi-channel functional near infrared spectroscopy (fNIRS), that would enable us to pinpoint specific brain regions and activation patterns in response to specific business decision making. This, in turn, would provide an empirical foundation for future development of a neuroeconomics model for predicting economic decision making in business contexts.

Multi-channel fNIRS is a portable and non-invasive imaging technique that measures cortical hemodynamic activities in the human brain. It quantifies the cerebral concentration changes of oxygenated hemoglobin (Δ[HbO]) and deoxygenated hemoglobin (Δ[Hb]) with high temporal resolution based on the alteration of optical absorption and scattering of near infrared (NIR) light propagating through the human brain. The temporal and spatial features of Δ[HbO] and Δ[Hb] are then used as biomarkers of neuronal activations^4,5^. In the last two decades, fNIRS has gained popularity and is recognized as a non-invasive tool to functionally image brain activations and/or diagnose brain diseases ^6,7^. Compared to fMRI, fNIRS is cost-effective, less sensitive to motion artifacts, and with higher temporal resolution, all of which makes it easier to use in a task-oriented, more-naturistic experimental environment.

In this study, we utilized the newsvendor problem (NP) for our experimental design. NP is one of the significant concepts associated with management science^8^; it refers to an prevalent business decision-making scenario, where an individual has to balance between potential loss and waste, in order to achieve maximum expected profit. The typical scenario is that of a store manager deciding the number of units of products to stock when the number of customers is uncertain^9^. Too little stock leads to potential loss sales; too much stock leads to potential waste. Risky decisions are needed to stock the inventory for profit under arbitrary requirements. In such decision contained risks, the likelihood of the consequence is known. However, the safe or risky outcome differs in terms of the reward ^10^. There is very little empirical research on the brain activities underlying decisions made under varying conditions of risk. This study aims to fill this void.

Specifically, the purpose of the study was to map and examine brain activities while subjects were making decisions required of the NP task. Previous neuroeconomics literature reported that there are three foremost risk decision-making processes, namely, reward processing, cognitive control, and social cognition ^11^. These processes trigger mainly dorsolateral prefrontal cortex (DLPFC), orbitofrontal cortex (OFC) and angular cingulate cortex (ACC) along with several other regions in the brain as a network shown in the fMRI studies ^12^. In this study, we hypothesized that the NP would stimulate both DLPFC and OFC significantly in the human brain, and that more challenging NPs would result in deactivation of DLPFC. These two hypotheses were based on prior findings that DLPFC represents cognitive control and emotion, and OFC is part of the reward pathway, both of which may be stimulated or activated at the cortical level during decision making under risks. To test our hypotheses, we designed and incorporated the NP using a computer game-based platform with simultaneous 77-channel fNIRS data acquisition.

## Results

Three statistical analyses were performed to test: (1) whether DLPFC and OFC were significantly evoked by the NP with respect to baseline regardless of treatment (HM or LM) types; (2) whether DLPFC and/or OFC were activated by HM or LM versus baseline; and (3) whether a more challenging NP task created significant difference in brain activation or deactivation in DLPFC or OFC regions.

First, Fig. 1(a) shows the front view of a topographic beta map (i.e., Δ[HbO]) evoked by the 40-trial NP from all 27 human subjects regardless of either HM or LM treatment. Since this study focused on NP-induced hemodynamic changes mainly in the OFC and DLPFC, we formed four clusters - I, II, III, and IV - based on several key Brodmann Areas, as shown by the four colored regions on the left hemisphere in Fig. 1(b). Based on the output of NIRS_SPM^13^, specifically, cluster I covers BAs 6 and 8, cluster II covers BAs 9 and 46 (i.e., DLPFC), cluster III covers BAs 10 and 11 (i.e., OFC), and cluster IV covers BAs 44 and 45 (i.e., Broca’s area). Fig. 1(c) is the corresponding t-map after one-sample t-tests, illustrating that the NP strongly activated clusters II, III, and IV but deactivated cluster I on the left hemisphere. On the other hand, the NP activated clusters III and IV and deactivated in cluster II on the right hemisphere. For more quantitative comparison, beta values of four clusters were calculated and plotted in Figs. 1(d) and 1(e) for both activation and deactivation, respectively. It is very clear that NP decision making significantly activated OFC on both hemispheres and DLPFC on the left hemisphere.

**Fig. 1.**
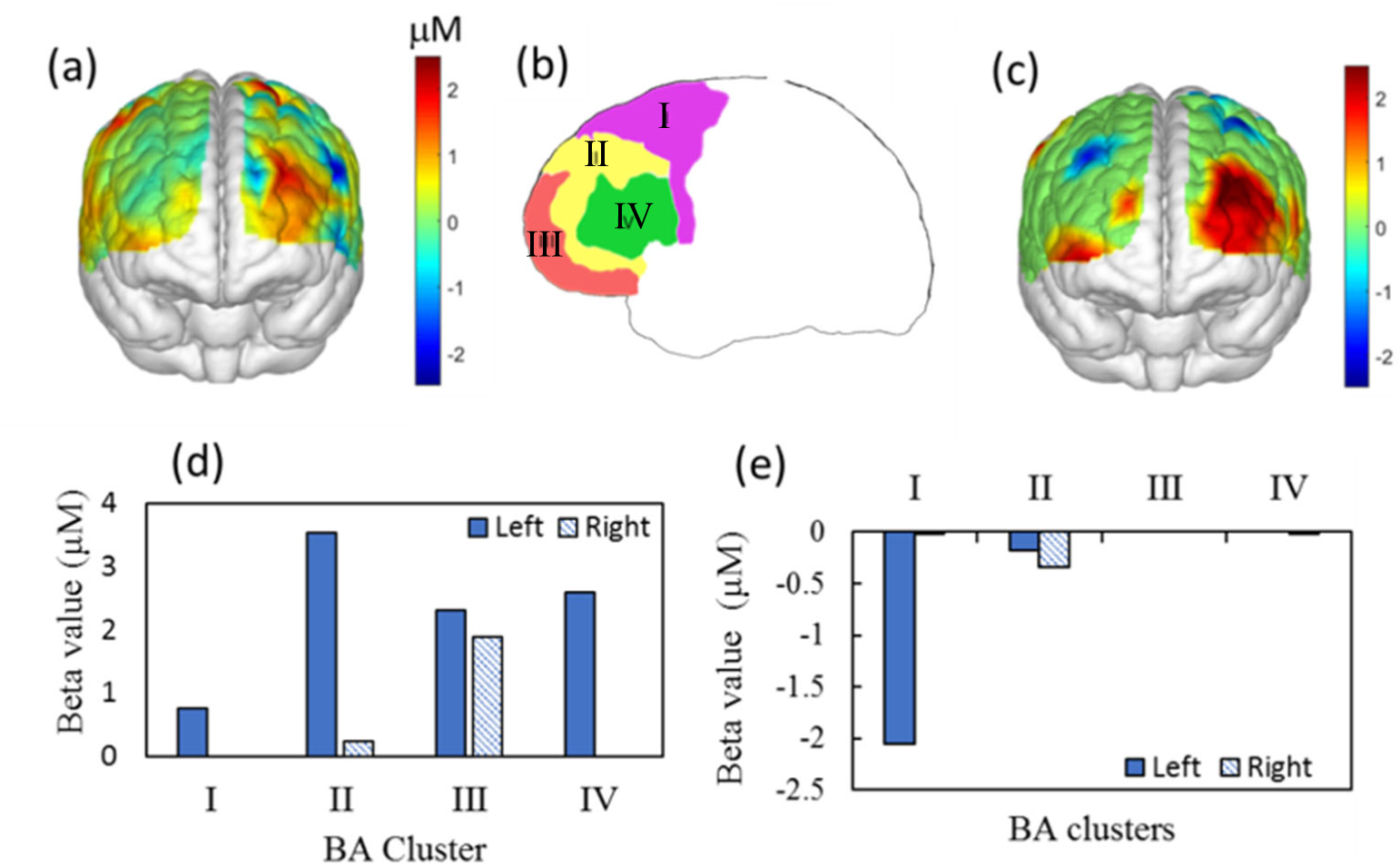
(a) It shows a topographic beta map stimulated by the 40-trial NP from all subjects (n=27) without considering the treatment level. Four dashed areas mark four BA clusters on the left hemisphere, as marked (b) clusters I, II, III, IV. Cluster I: BAs 6 and 8; cluster II: BAs 9 and 46 (i.e., DLPFC); cluster III: BAs 10 and 11 (i.e., OFC); cluster IV: BAs 44 and 45 (i.e., Broca’s area). (c) The corresponding t-map compared to the zero baseline. NP-evoked (d) activation and (e) deactivation beta values in respective BAs.

Second, following the same format as Figs. 1(c) to 1(e), Figs. 2(a) – 2(c) show a front-view t-map with respect to the zero baseline, activated and deactivated beta values of each cluster induced by LM treatment, respectively. This set of figures demonstrate unambiguously that the NP decision making with LM did not evoke the OFC (i.e., cluster III), but rather activated strongly the left DLPFC and Broca’s area (i.e., clusters II and IV) and deactivated the right DLPFC (r-DLPFC, cluster II). On the other hand, Figs. 2(d) – 2(f) show similar figures of t-map, activated and deactivated beta values induced by HM treatment, respectively. This set of figures evidently illustrate that the NP decision making with HM treatment significantly activated OFC (cluster III) on both hemispheres and left DLPFC (l-DLPEC).

**Fig. 2.**
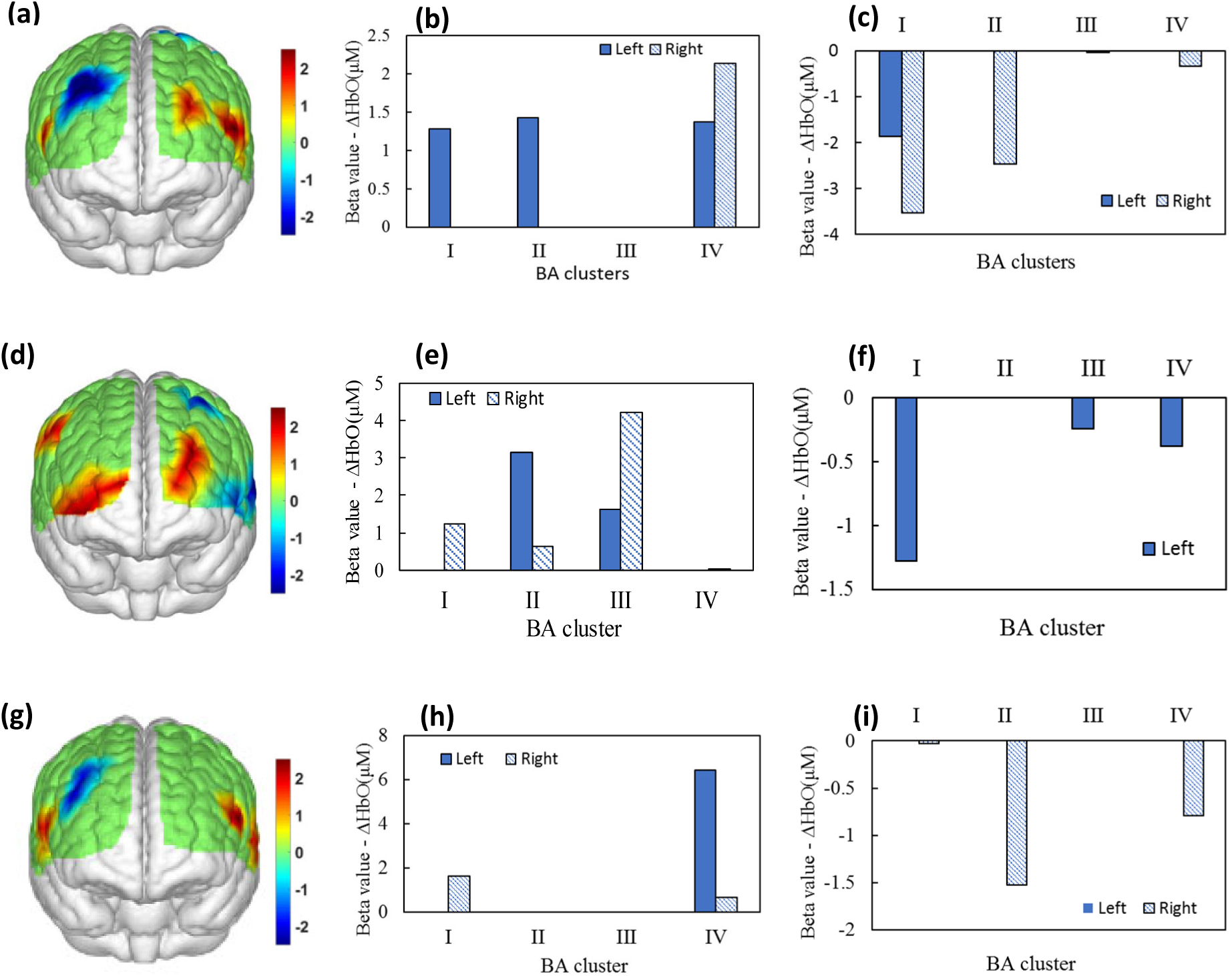
Front-view topographic t-maps of Δ[HbO] evoked by the NP with (a) LM treatment (n=14) and (d) HM treatment (n=13). One-sample t-tests were performed for these two cases. (b) and (c) represent the beta values with respect to clusters under LM. (e) and (f) mark the beta values with respect to clusters under HM. (g) It shows a front-view topographic t-map comparing activation/deactivation between LM and HM tasks based on two-sample t-tests. (h) and (i) mark the differential beta values between LM and HM treatment for each cluster.

Third, Fig. 2(g) shows a topographic t-map obtained by two-sample t-tests for comparing NP-evoked brain activation or deactivation under LM and HM treatment. Accordingly, Figs. 2(h) and 2(i) plot differential beta values between the two treatment cases. Two major observational features are noted: A more challenging NP with LM created significant deactivation in r-DLPFC and significant activation in left cluster IV.

## Discussion

In a complex and turbulent business environment, managers have to make decisions that are fraught with risks. Risky decision making is a complex cognitive process, requiring contribution and integration of actions from multiple regions of the human brain. The scope of this study was to observe which regions of the human brain are responsible for making risky decisions in a business context, as represented by the widely studied newsvendor problem (NP). The hypotheses were that NP stimulates both DLPFC and OFC significantly in the human brain, and that more challenging NP with LM treatment triggers more deactivation in DLPFC than HM treatment, based on 77-channel fNIRS measurements from 27 human subjects.

First, our results (Fig. 1) confirmed that both DLPFC and OFC regions, particularly the left side of DLPFC, played significant and key roles in cognitive processing when subjects had to solve the NP regardless of difficulty levels. Several previous studies in the literature have reported that DLPFC is responsible for risky decision making (in non-business context) and cognitive control in terms of planning and working memory ^5, 12,14,15^. Our findings in this paper are consistent with those reported in these documents. Furthermore, some parts of Broca’s area (cluster IV) was also activated during the tasks. It is known that Broca’s area is primarily responsible for language processing^16^ and speech, both outer and inner speech^17^. In some scenarios, it interacts with the inferior frontal gyrus to process language information for those who use second languages^16^. Evidence of this interaction was found in our study, possibly because most of our subjects were international students who relied on language processing during the decision making tasks^18^. Furthermore, some studies have shown that Broca’s area along with inferior frontal gyrus is responsible for self-talk. This scenario has been reported prominent when self-encouragement is needed to gain success, which are seen in sports literature^19,20^. In the case of LM trials, there was a high probability to lose profits if subjects did not make a careful decision. Once they lose, self-encouragement may occur for continuing to play the game for profits, leading to strong activation in Broca’s area.

Second, Left DLPFC is more prominently activated in the high margin case where it is easier to create a goal-oriented decision with a less risky environment. In a fMRI-based study where the subjects were given the task of “Tower of London”, a popular protocol based on planning, it was observed that during the goal hierarchical state the left DLPFC was significantly activated^21^.

Another study reported significant DLPFC stimulations when the subjects made decision to execute a right-hand movement to enter the order quantity on the computer keyboard^22^. Furthermore, we observed that OFC was activated during the HM NP task. Since OFC is suggested to be one of the significant components in the reward pathway^23^, the activation of OFC explains the anticipation for gains during the trials. Third, compared to HM stimulation, several distinct features of brain responses to LM stimulation are noted: (1) strong activation in left Broca’s area, (2) significant deactivation in r-DLPFC, and (3) no activation/deactivation in OFC. Since this set of tasks were more challenging, subjects needed to pay more attention to the tasks and to control their emotions during the tasks. Since BA 45 (i.e., Broca’s area) has a function of modulating emotional response besides language process, it is reasonable to observe strong activation in this cortical area in response to the LM tasks. The reason for strong r-DLPFC deactivation may be attributed to intense stress due to more difficult decision-making challenges^24,25^. Although OFC is expected to have strong response in this case because it plays a key role in decision making involving reward, the observation in our study was beyond our expectation. A plausible reason for this finding is that, as the more challenging NP tasks shifted the subjects’ attention from a less-stress reward phase to a more-stress defying phase, the brain activations occurred in DLPFC rather than in OFC. It is known that DLPFC promotes decision-making process, pre-motor cortex, and supplementary motor cortex, which are responsible for planning execution of movements^26^.

Finally, to be more rigorous in testing for significant differences in brain responses to LM and HM treatment, two-sample t-tests were performed. The results confirmed that LM tasks triggered (1) significant activation in Broca’s area (i.e., left cluster IV) and (2) significant deactivation in r-DLPFC. The reason for the former observation was mentioned above, namely, Broca’s area has a function of modulating emotional response. The latter observation is consistent with results reported in a few recent publications. For example, an fMRI study involving 27 subjects suggested that deactivation in r-DLPFC may occur due to acute stress which weakens high-level cognitive functions such as working memory^27^. In addition, a transcranial direct current stimulation (tDCS) based study with 120 participants showed that tDCS delivered on the r-DLPFC can prevent stress-induced working memory deficits ^24^. In our study, the NP with the LM protocol was rather risky and a bit lengthy, subjecting the participants to higher levels of stress. Consequently, we observed clear and significant deactivation in r-DLPEC only during LM tasks.

### Summary and future work

In summary, this study showed that NP-based decision-making tasks stimulated key brain areas such as DLPFC and OFC for high-level cognitive functions based on 77-channel simultaneous hemodynamic measurements from 27 human control subjects. The study observed that there were multiple regions activated and deactivated as responses to the tasks. Specifically, DLPFC and OFC were significantly evoked by NP tasks versus baseline regardless of treatment types. DLPFC and OFC were activated by HM versus baseline, while l-DLPFC and Broca’s area were strongly activated during LM versus baseline. Furthermore, significant deactivation in r-DLPFC was observed and attributed to challenging stress created by the LM with respect to HM. Overall, this study proved our hypotheses that DLPFC and OFC are two key cortical regions when performing NP tasks, revealing that right DLPFC in particular gets more deactivated under challenging risky decision making.

Since this study consisted of only 27 subjects, separated into two different groups for LM and HM treatments, the sample size was statistically low. Therefore, it is appropriate for future studies to corroborate the findings of this study using larger sample sizes.

## Material and Methods

### Participants

A total of 27 subjects (20 males and 7 females) with a mean of 23 ± 5 years of age participated in the study. They were randomly assigned into two experimental groups with different (i.e., HM or LM) treatment (risk) levels. The inclusion criteria included: either sex, any ethnic background, and in an age range of 18−40 years old. The exclusion criteria included: (1) diagnosed with a psychiatric disorder, (2) history of a neurological condition, or severe brain injury, or violent behavior, (3) prior institutionalization or imprisonment, and (4) current intake of any medicine or drug. Specifically,13 of them attended a high-profit margin treatment, while the other 14 attended a low-profit margin treatment. The study protocol was approved by the institutional review board (IRB) of the University of Texas at Arlington and complied with all applicable federal guidelines. Informed consent was obtained from each participant prior to each experiment.

### NP Protocol Design

This study design was based on the Newsvendor Problem^8^, where a news vendor must decide how many newspapers to buy each day at the wholesale price and to sell them at the retail price. This problem has five major characteristics: (1) the demands are uncertain but from a known distribution; (2) the decision must be taken for every period; (3) there is a cost for ordering too many items; (4) the number of items ordered at each time is called *order quantity*, which must be decided for the inventory by the subject in each trial; (5) each trial is independent. According to the NP model^7,9,28^, under the condition that order quantity is larger than the unknown demand, the final profit (π) can be calculated as a function of order quantity and unknown demand by Equation 1^28^.

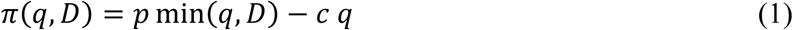

where *q* denotes the order quantity, and *D* represents the unknown demand. *c* is the cost, and *p* is the price to sell.

Based on the theory above, the experimental protocol was designed with two independent treatments, namely, the high-profit margin (*c* ≪ *p*) and low-profit margin (*c* is close to *p*). Table 1 shows the information of the design for two different treatments. In the high-profit margin (HM) treatment, the price to sell, *p*, was designed to be $32 while the cost was only $8, all of which resulted in lower risk of losing profits. On the other hand, in the low-profit margin (LM) treatment, while keeping *p* still $32, the cost was raised to $24, leading to higher risk of losing profits. The demand was kept unknown until the participant made his or her decision by typing the order quantity between 0 to 300 per trial. Then, the given demand was randomly generated from a uniform distribution between 0 to 300. The history of demands from previous trials were visible to the participant, based on which the participant could decide for the current trial. There were 40 trials in total in each experiment. The selling price and cost were kept constant and known for each HM and LM treatment.

**Table 1:** NP Protocol Summary.

| Treatment | High-profit Margin (HM) | Low-profit Margin (LM) |
| --- | --- | --- |
| Price ($p$) | \$32 | \$32 |
| Cost ($c$) | \$8 | \$24 |
| Demand Distribution | Uniform [0 300] | Uniform [0 300] |

### NP Protocol Implementation

One entire experiment consisted of a 30-s baseline and 40 blocks of NP tasks, as shown in Fig. 3. The 30-s ‘baseline’ was needed to acquire the baseline of cerebral hemodynamic functions of each subject. Each block contained one NP trial, and each NP trial had 4 phases: decision, rest, feedback, and rest. The variable ‘decision’ phase lasted for a maximum of 20 seconds, during which the subject was asked to make decisions on order quantity given such visible information as price, cost, and demand distribution range for either HM or LM group. The subject had to enter the quantity in a text box on the screen within the 20-s maximal time window.

**Fig. 3.**
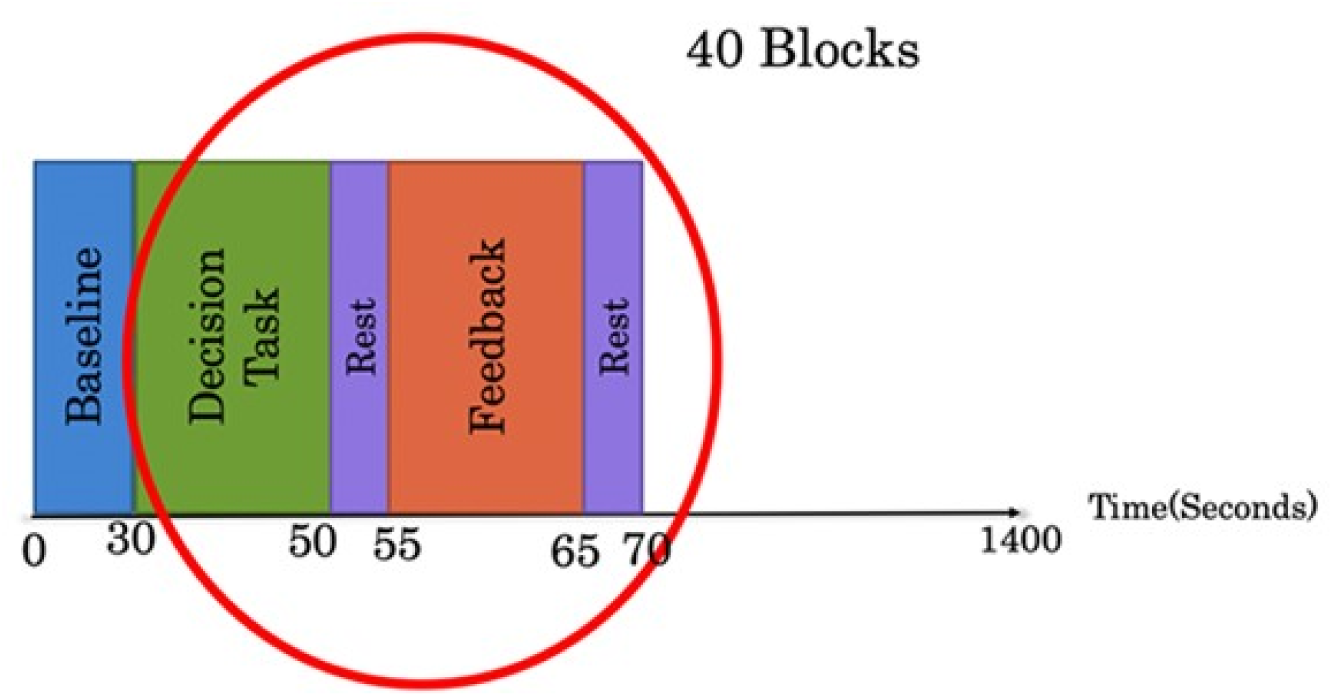
The NP experimental protocol consisting of a 30-s initial baseline and 40 blocks. Each block (marked by the red circle) has 4 phases: up to 20-s NP Decision Task, 5-s Rest, 10-s Feedback, and 5-s Rest again before starting another block. Each block lasts about 30-40 seconds.

After this phase, the screen shifted to a 5-s ‘rest’ phase, followed by the 10-s ‘feedback’ phase. Within this 10-s period, the subject was shown a summary table listing the price, cost, demand distribution, profit/loss and the cumulative profits/losses that were made after every trial. Then the protocol proceeded to the next trial following another 5-s ‘rest’ period. All the profit/loss details were stored along with the corresponding time stamps.

### fNIRS Experiments

A continuous-wave, multi-channel fNIRS system (LABNIRS, Shimadzu Corp., Kyoto, Japan) was employed in this study. As reported before ^29^, a customized 77-channel layout incorporating 25 pairs of laser transmitters and 23 light receivers was used to cover from the prefrontal cortex to sensorimotor cortex. Figure 4 shows the locations of 77 channels on a human brain template; these channels were connected to the LABNIRS data collection system. The distance between the nearest source and detector fiber optodes was 3 cm, resulting in a sensitive detection depth of 1.5-2 cm under the scalp. All the source and detector fibers were firmly and steadily held by the whole-head helmet on each subject’s head; the data sampling frequency was 12.82 Hz. Spatial co-registration measurements were taken for each subject after each experiment by a 3D digitizer (FASTRACK, Polhemus, Colchester, VT, USA).

**Fig. 4.**
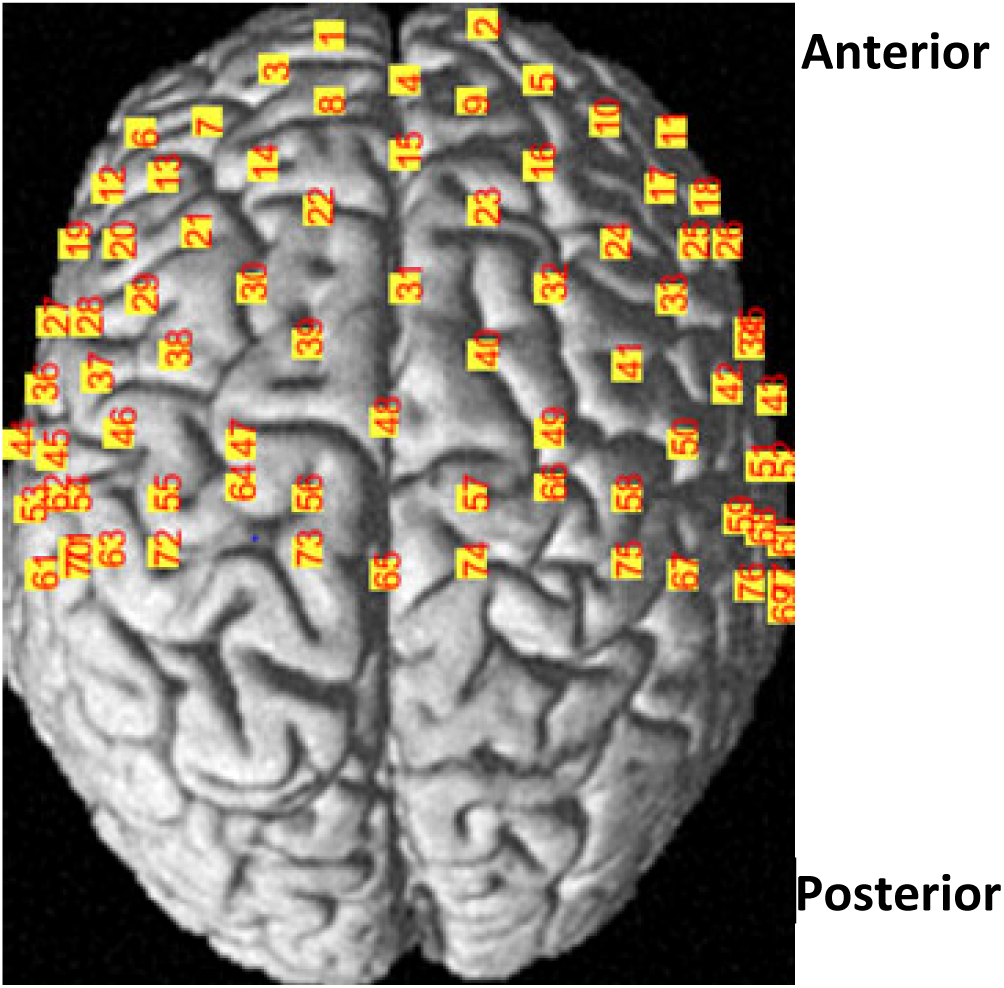
A channel-wise layout of fNIRS optodes on a human brain template, with labels of 77 channels derived from 25 pairs of light sources and 23 detectors.

### Data Preprocessing

The raw outputs of the fNIRS system were time-dependent optical intensities at three wavelengths (i.e. 780 nm, 805 nm, and 830 nm), the alterations of which were affected by the changes of hemoglobin concentrations. A band-pass filter of 0.01 – 0.2 Hz was applied to remove artifacts from cardiac pulses (∼0.8-1.2 Hz), respiration (∼ 0.2 - 0.3 Hz)^30^, muscle movements, and systemic drifts. Next, the modified Beer-Lambert Law was used to convert the recorded optical intensity at 780 nm and 830 nm into relative oxygenated hemoglobin concentration, Δ[HbO], and relative de-oxygenated hemoglobin(Δ[Hb])^4,31^. Then, each time series of Δ[HbO] and Δ[Hb] was baseline calibrated by subtracting its temporal average to remove any potential drifts during the baseline recording time. Moreover, to further remove physiological variations due to arterial blood pressure or Mayer waves coming from the scalp and skull, we subtracted the signal spatially averaged across all 77 channels to remove the global noise ^29,32,33^.

### Activation Maps Based on the General Linear Model

For each channel, a preprocessed time series of Δ[HbO] was used as the input for the general linear model (GLM) analysis using MATLAB. In the process, the design temporal matrix was generated using a boxcar function reflecting the NP task blocks convoluted with a canonical hemodynamic response function (HRF) ^16,34^. For each NP experiment (with HM or LM treatment), two regressors were generated corresponding to the decision phase and feedback phase, while the data analysis was mainly focused on the decision phase. The activation amplitudes (i.e., beta values) for both regressors were calculated or fitted through a regression algorithm between calculated Δ[HbO] time series and the design temporal matrix using a weighted least square method ^35^. Such regression processes were carried out for each channel, giving rise to an array of 77 fitted beta values relative to their own baselines for each human subject.

Next, statistical analysis using one-sample or two-sample t-tests was performed to test our hypotheses. Specifically, we wished to know (1) whether DLPFC and/or OFC were significantly evoked by NP versus baseline regardless of treatment (HM or LM) types, and (2) whether DLPFC and/or OFC were activated by HM or by LM versus baseline. Furthermore, a key question was whether a more challenging NP created significant difference in brain activation or deactivation in DLPFC or OFC regions. For test (1), one-sample t-tests (against the mean of zero baseline) were performed over beta values of Δ[HbO] from all 27 subjects for each of the 77 channels, without considering any treatment effect. This step allowed us to map/identify significant cerebral activation/deactivation regions (i.e., channels) while the subjects made risky NP decisions. For test (2), beta values of Δ[HbO] were analyzed independently for subjects with either LM or HM treatment, respectively. Similar one-sample t-tests were performed among 13 participants with HM treatment and separately among 14 participants with LM, at each channel to examine the statistical significance versus the mean of zero baseline. For the final test, two-sample t-tests were performed at each channel between the two groups (with HM and LM treatments) to determine significant differences in brain activation/deactivation caused by the NP. Accordingly, topographic t-value maps were generated using easytopo 2.0 ^36^.

### Beta Value Extraction with respect to Broadman Areas

We identified several key Brodmann areas (BAs): BAs 6, 8, 9, 10, 11, 44, 45, and 46 to mark or co-register NP-evoked cortical regions sensed/detected by fNIRS channels on each participant’s cortex. The underlying neuroanatomy of those areas are marked as follows: BA 6 corresponds to the premotor cortex, BA 8 corresponds to the frontal eye fields, BAs 10, 11, and 47 to OFC, BAs 9 and 46 to DLPFC, and BAs 44 and 45 to Broca’s area. Several steps of BAs-based analysis were taken to quantify beta values of HbO changes in each respective BA. First, a software package, named NIRS_SPM^13^, was used not only to co-register between the selected BAs and each channel location (see Fig. 4) but also to quantify activation values of ΔHbO from each channel and corresponding percentage of overlap between the optical detection area and respective anatomical regions or BAs, as demonstrated in Fig. 5. Second, the beta value of each channel determined by the GLM algorithm was multiplied by the percentage of overlap, as a weighting factor, for each of the corresponding BAs. Next, the beta value of each key BA on each hemisphere was estimated by summing up weighted beta values from all channels that covered each of respective BAs. Finally, both activations and deactivations in each BA were determined and plotted for each of the key BAs.

**Fig.5.**
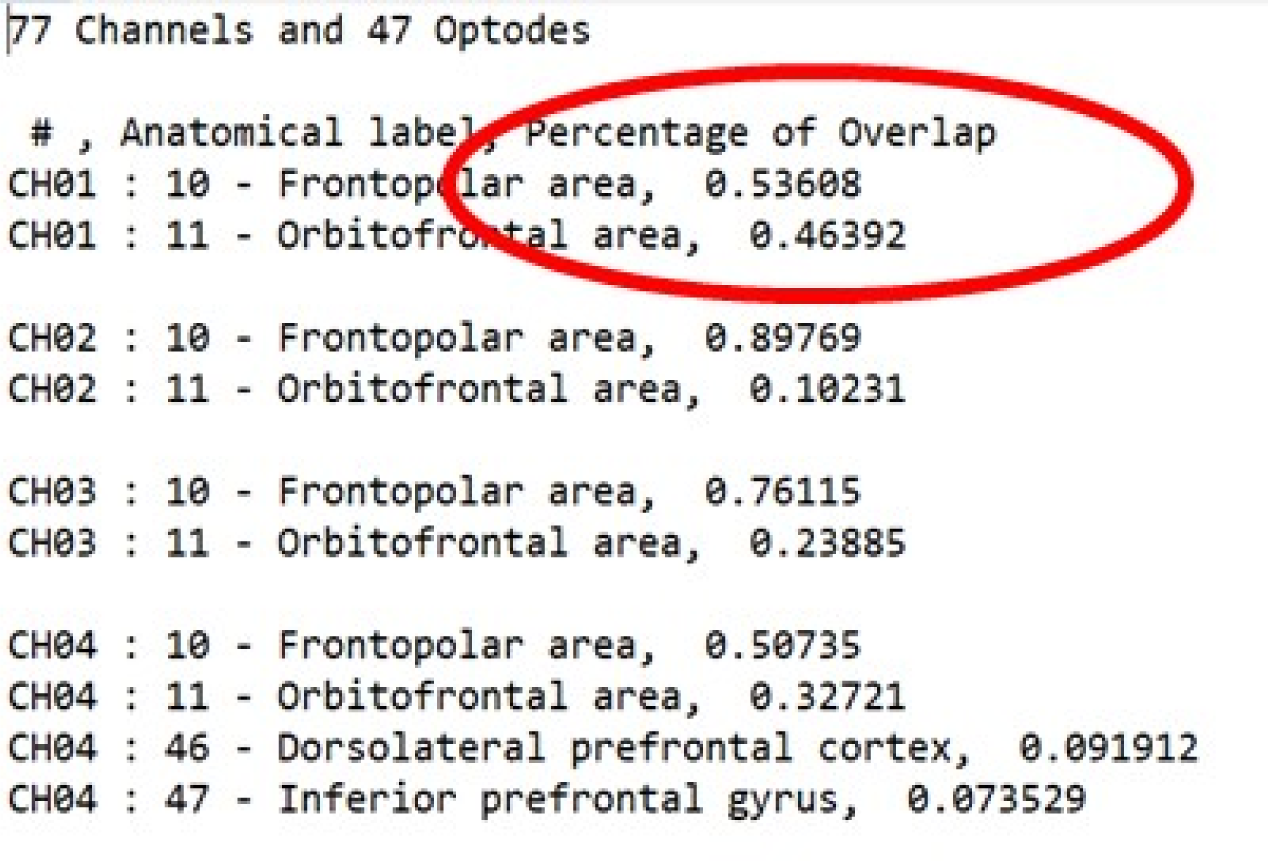
Relationship between optical channels (e.g. CH01), anatomical labels (e.g., frontopolar area, and percentage of overlap (e.g. 0.53608) between the optical detection area by optical channels and respective anatomical regions or BAs.

## Acknowledgments

The authors acknowledge the support in part from the seed funding from the University of Texas at Arlington.

## Author contributions

H.W. performed the experiment, analyzed the data, interpreted the results, and prepared the manuscript. Y. L. recruited human participants, implemented the Newsvendor experiment protocols, and coordinated the experiment. X. W. analyzed the data, discussed the results, and participated in manuscript revision. S. N. and K-Y C. designed the Newsvendor experiment, discussed the results, and revised the manuscript. H. L. initiated and supervised the study, discussed and interpreted the results, as well as reviewed and revised the manuscript.

## References

1 Hsu, M., Bhatt, M., Adolphs, R., Tranel, D. & Camerer, C. F. Neural systems responding to degrees of uncertainty in human decision-making. Science 310, 1680–1683, doi: 10.1126/science.1115327 (2005).

2 Xue, G., Chen, C., Lu, Z. L. & Dong, Q. Brain Imaging Techniques and Their Applications in Decision-Making Research. Xin Li Xue Bao 42, 120–137, doi: 10.3724/SP.J.1041.2010.00120 (2010).

3 Glimcher, P. W., Camerer, C., Poldrack, R. A. & Fehr, E. Neuroeconomics: Decision Making and the Brain. (Oxford: Academic Press, 2008).

4 Boas, D. A., Dale, A. M. & Franceschini, M. A. Diffuse optical imaging of brain activation: approaches to optimizing image sensitivity, resolution, and accuracy. NeuroImage 23 Suppl 1, S275–288, doi: 10.1016/j.neuroimage.2004.07.011 (2004).

5 Cazzell, M., Li, L., Lin, Z. J., Patel, S. J. & Liu, H. Comparison of neural correlates of risk decision making between genders: an exploratory fNIRS study of the Balloon Analogue Risk Task (BART). NeuroImage 62, 1896–1911, doi: 10.1016/j.neuroimage.2012.05.030 (2012).

6 Boas, D. A., Elwell, C. E., Ferrari, M. & Taga, G. Twenty years of functional near-infrared spectroscopy: introduction for the special issue. NeuroImage 85 Pt 1, 1–5, doi: 10.1016/j.neuroimage.2013.11.033 (2014).

7 Benzion, U., Cohen, Y., Peled, R. & Shavit, T. Decision-Making and the Newsvendor Problem: An Experimental Study. Journal of the Operational Research Society 59, 1281–1287, doi: 10.1057/palgrave.jors.2602470 (2008).

8 Cohen, Y., Peled, R. & Shavit, T. Decision-making and the newsvendor problem: an experimental study. J. Oper. Res. Soc. 59, 1281–1287 (2008).

9 Gaspars-Wieloch, H. Newsvendor Problem under Complete Uncertainty: A Case of Innovative Products. Central European Journal of Operations Research 25, 561–585, doi: 10.1007/s10100-016-0458-3 (2017).

10 Krain, A. L., Wilson, A. M., Arbuckle, R., Castellanos, F. X. & Milham, M. P. Distinct Neural Mechanisms of Risk and Ambiguity: A Meta-Analysis of Decision-Making. NeuroImage 32, 477–484, doi: 10.1016/j.neuroimage.2006.02.047 (2006).

11 Declerck, C. & Boone, C. Neuroeconomics of Prosocial Behavior: The Compassionate Egoist. (Academic Press, 2015).

12 Farrar, D. C., Mian, A. Z., Budson, A. E., Moss, M. B. & Killiany, R. J. Functional Brain Networks Involved in Decision-Making under Certain and Uncertain Conditions. Neuroradiology 60, 61–69, doi: 10.1007/s00234-017-1949-1 (2018).

13 Ye, J. C., Tak, S., Jang, K. E., Jung, J. & Jang, J. NIRS-SPM: statistical parametric mapping for near-infrared spectroscopy. NeuroImage 44, 428–447, doi: 10.1016/j.neuroimage.2008.08.036 (2009).

14 Makwana, A. & Hare, T. Stop and be fair: DLPFC development contributes to social decision making. Neuron 73, 859–861, doi: 10.1016/j.neuron.2012.02.010 (2012).

15 Huang, D. et al. Activation of the DLPFC Reveals an Asymmetric Effect in Risky Decision Making: Evidence from a tDCS Study. Front Psychol 8, 38, doi: 10.3389/fpsyg.2017.00038 (2017).

16 Buchsbaum, B. R., Hickok, G. & Humphries, C. Role of Left Posterior Superior Temporal Gyrus in Phonological Processing for Speech Perception and Production. Cognitive Science 25, 663–678, doi: 10.1207/s15516709cog2505_2 (2001).

17 Morin, A. Self-Awareness Part 2: Neuroanatomy and Importance of Inner Speech. Social and Personality Psychology Compass 5, 1004–1017, doi: 10.1111/j.1751-9004.2011.00410.x (2011).

18 Kim, K. H. S., Relkin, N. R., Lee, K.-M. & Hirsch, J. Distinct Cortical Areas Associated with Native and Second Languages. Nature 388, 171–174, doi: 10.1038/40623 (1997).

19 St Clair Gibson, A. & Foster, C. The role of self-talk in the awareness of physiological state and physical performance. Sports Med 37, 1029–1044, doi: 10.2165/00007256-200737120-00003 (2007).

20 Mitchell, R. W. Self-awareness without inner speech: a commentary on Morin. Conscious Cogn 18, 532–534, doi: 10.1016/j.concog.2008.12.003 (2009).

21 Kaller, C. P., Rahm, B., Spreer, J., Weiller, C. & Unterrainer, J. M. Dissociable Contributions of Left and Right Dorsolateral Prefrontal Cortex in Planning. Cerebral Cortex (New York, N.Y.: 1991) 21, 307–317, doi: 10.1093/cercor/bhq096 (2011).

22 Hoshi, E. & Tanji, J. Area-Selective Neuronal Activity in the Dorsolateral Prefrontal Cortex for Information Retrieval and Action Planning. Journal of Neurophysiology 91, 2707–2722, doi: 10.1152/jn.00904.2003 (2004).

23 Bolla, K. I. et al. Orbitofrontal Cortex Dysfunction in Abstinent Cocaine Abusers Performing a Decision-Making Task. NeuroImage 19, 1085–1094, doi: 10.1016/S1053-8119(03)00113-7 (2003).

24 Bogdanov, M. & Schwabe, L. Transcranial Stimulation of the Dorsolateral Prefrontal Cortex Prevents Stress-Induced Working Memory Deficits. J. Neurosci. 36, 1429–1437, doi: 10.1523/JNEUROSCI.3687-15.2016 (2016).

25 Qin, S., Hermans, E. J., van Marle, H. J. F., Luo, J. & Fernández, G. Acute Psychological Stress Reduces Working Memory-Related Activity in the Dorsolateral Prefrontal Cortex. Biological Psychiatry 66, 25–32, doi: 10.1016/j.biopsych.2009.03.006 (2009).

26 Schubotz, R. I. & von Cramon, D. Y. Functional–Anatomical Concepts of Human Premotor Cortex: Evidence from fMRI and PET Studies. NeuroImage 20, S120–S131, doi: 10.1016/j.neuroimage.2003.09.014 (2003).

27 Dias, E. C. & Segraves, M. A. Muscimol-induced inactivation of monkey frontal eye field: effects on visually and memory-guided saccades. J Neurophysiol 81, 2191–2214, doi: 10.1152/jn.1999.81.5.2191 (1999).

28 Schweitzer, M. E. & Cachon, G. P. Decision Bias in the Newsvendor Problem with a Known Demand Distribution: Experimental Evidence. Management Science 46, 404–420, doi: 10.1287/mnsc.46.3.404.12070 (2000).

29 Cacola, P., Getchell, N., Srinivasan, D., Alexandrakis, G. & Liu, H. Cortical activity in fine-motor tasks in children with Developmental Coordination Disorder: A preliminary fNIRS study. Int J Dev Neurosci 65, 83–90, doi: 10.1016/j.ijdevneu.2017.11.001 (2018).

30 Erdoğan, S. B., Yücel, M. A. & Akin, A. Analysis of Task-Evoked Systemic Interference in fNIRS Measurements: Insights from fMRI. NeuroImage 87, 490–504, doi: 10.1016/j.neuroimage.2013.10.024 (2014).

31 Nguyen, T. et al. Exploring brain functional connectivity in rest and sleep states: a fNIRS study. Sci Rep 8, 16144, doi: 10.1038/s41598-018-33439-2 (2018).

32 Desjardins, A. E., Kiehl, K. A. & Liddle, P. F. Removal of Confounding Effects of Global Signal in Functional MRI Analyses. NeuroImage 13, 751–758, doi: 10.1006/nimg.2000.0719 (2001).

33 Tak, S. & Ye, J. C. Statistical Analysis of fNIRS Data: A Comprehensive Review. NeuroImage 85, 72–91, doi: 10.1016/j.neuroimage.2013.06.016 (2014).

34 Schroeter, M. L. et al. Towards a Standard Analysis for Functional Near-Infrared Imaging. NeuroImage 21, 283–290, doi: 10.1016/j.neuroimage.2003.09.054 (2004).

35 Tian, F. et al. Prefrontal responses to digit span memory phases in patients with post-traumatic stress disorder (PTSD): A functional near infrared spectroscopy study. NeuroImage. Clinical 4, 808–819, doi: 10.1016/j.nicl.2014.05.005 (2014).

36 Tian, F. et al. Test-retest assessment of cortical activation induced by repetitive transcranial magnetic stimulation with brain atlas-guided optical topography. Journal of biomedical optics 17, 116020, doi: 10.1117/1.JBO.17.11.116020 (2012).

